# scMM: Mixture-of-experts multimodal deep generative model for single-cell multiomics data analysis

**DOI:** 10.1101/2021.02.18.431907

**Authors:** Kodai Minoura, Ko Abe, Hyunha Nam, Hiroyoshi Nishikawa, Teppei Shimamura

## Abstract

The recent development in single-cell multiomics analysis has enabled simultaneous detection of multiple traits at the single-cell level, thus providing deeper insights into the cellular phenotypes and functions in diverse tissues. However, currently, it is challenging to infer the joint representations and learn relationships among multiple modalities from complex multimodal single-cell data. Herein, we present scMM, a novel deep generative model-based framework for the extraction of interpretable joint representations and cross-modal generation. scMM addresses the complexity of data by leveraging a mixture-of-experts multimodal variational autoencoder. The pseudocell generation strategy of scMM compensates for the limited interpretability of deep learning models and discovered multimodal regulatory programs associated with latent dimensions. Analysis of recently produced datasets validated that scMM facilitates high-resolution clustering with rich interpretability. Furthermore, we show that cross-modal generation by scMM leads to more precise prediction and data integration compared with the state-of-the-art and conventional approaches.

## 1 Introduction

Recent technological advances have enabled simultaneous acquisitions of multiple omics data at the resolution of a single-cell, thus producing “multimodal” single-cell data [1, 2, 3, 4, 5, 6]. These technologies offer additional measurements, such as immunophenotypes or chromatin accessibility in conjunction with transcriptome information. Research studies using emerging multimodal single-cell technologies have contributed to exciting, biologically important discoveries in various fields, including immune cell profile and cell fate decision that could not have been elucidated with the use of only one modality [5, 6].

Conversely, some obstacles need to be overcome to computationally extract useful knowledge from highly complex, single-cell multimodal data. First, it is challenging to infer the low-dimensional joint representations from multiple modalities that can be used for downstream analyses such as clustering. Second, although multimodal single-cell data allow the learning of relationships among modalities that could be used to train prediction models, many-to-many predictions of single-cell data (e.g., from single-cell transcriptome to chromatin accessibility) with high accuracy is still an unsolved problem. These problems are mainly attributed to the difficulties associated with capturing latent common factors and relationships across modalities, which differ significantly in characteristics, including data distribution, dimensionality, and sparsity.

There are several methods recently developed for the analysis of single-cell multimodal data. Although they aim to address tasks such as latent feature extraction, their performance is currently limited in different aspects. Methods based on generalized linear models, such as Seurat and scAI, often fail to capture complex structures in single-cell data [5, 15]. One powerful approach to capture nonlinear latent structures is to use expressive variational autoencoders (VAEs), which consist of a pair of neural networks wherein one encodes data into the latent space, and the other decodes them to reconstruct the data distribution [7]. scMVAE and totalVI are the currently available VAE-based methods for single-cell multimodal data analysis [13, 14]. Nevertheless, scMVAE requires a simplified conversion of chromatin accessibility to transcriptome before training, which is known to lead to the nonnegligible loss of epigenetic information [15]. Additionally, these models suffer from the “black-box” nature of deep learning models, making the interpretations of latent variables difficult. Lastly, none of these VAE-based methods are designed for the predictions across modalities.

To address these limitations, we have developed scMM, a novel statistical framework for single-cell multiomics analysis specialized in interpretable joint representation inference and predictions across modalities. scMM is based on a mixture-of-experts multimodal deep generative model and achieves end-to-end learning by modeling raw count data in each modality based on different probability distributions. Using recently published datasets produced by cellular indexing of transcriptomes and epitopes with sequencing (CITE-seq) and simultaneous high-throughput assay for transposase-accessible chromatin (ATAC) and RNA expression with sequencing (SHARE-seq), we demonstrate that scMM effectively extracts biologically meaningful latent variables encoding multimodal information. We show that these latent variables enable high-resolution clustering to reveal cellular heterogeneity that was not discovered in the original report [5, 6]. By leveraging the generative nature of the model, scMM provides users with multimodal “regulatory programs” that are associated with latent dimensions, thus aiding the interpretation of the results. Finally, exploration of the cross-modal generation of single-cell data by scMM proved that it outperforms the state-of-the-art prediction tool and contributes to more accurate integration of single-cell data from different modalities.

## 2 Results

### 2.1 The scMM model

scMM takes multimodal single-cell data as input, which contains measurements for multiple modalities across each cell. Let *x*_*n,m*_ be the feature vector for *m*-th modality in cell *n*. Theoretically, m can be any arbitrary number, although this study primarily focuses on the dualomics analysis since most recently developed multiomics methods deal with information of two modalities. We modeled *x*_*n,m*_ with probability distributions that capture the characteristics of data distributions for each modality. For transcriptome and surface protein data, negative binomial (NB) distribution was selected to explain non-negative counts with overdispersion [14]. In addition, chromatin accessibility data is non-negative count data; however, it exhibits extreme sparsity due to low signal (only two locus exist for each diploid cell), limited coverage, and closed chromatin. Therefore, we chose the zero-inflated negative binomial (ZINB) distribution for chromatin accessibility data. Although transcriptome data also show high sparsity, recent reports showed NB distribution is sufficient to explain the abundance of zeros in the transcriptome data [16, 9]. In contrast to some recently developed probabilistic models for chromatin accessibility that involves binarization, scMM models raw peak counts, allowing the natural increase of peak counts with sequencing depth [10, 17].

A conceptual view of scMM is shown in Fig. 1. scMM model for dualomics analysis consists of four neural networks wherein a pair of an encoder and a decoder is present in each modality. Let ***z*** be the set of low-dimensional vectors (set here to 10 dimensions) of latent variables. Encoders are used to infer the variational posterior 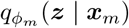 from which ***z***_*m*_ is sampled. Conversely, decoders calculate the parameters of negative binomial (NB) or zero-inflated negative binomial (ZINB) distributions, which can be written as 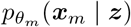. Herein, *ϕ*_*m*_ and *θ*_*m*_ denote the parameters for the encoder and decoder for the *m*-th modality, respectively. The scMM uses a MoE to factorize the joint variational posterior (see Methods). Accordingly, multimodal latent variables that encode information on two modalities can be obtained from MoE: 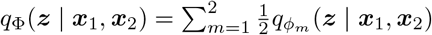.

**Figure 1:**
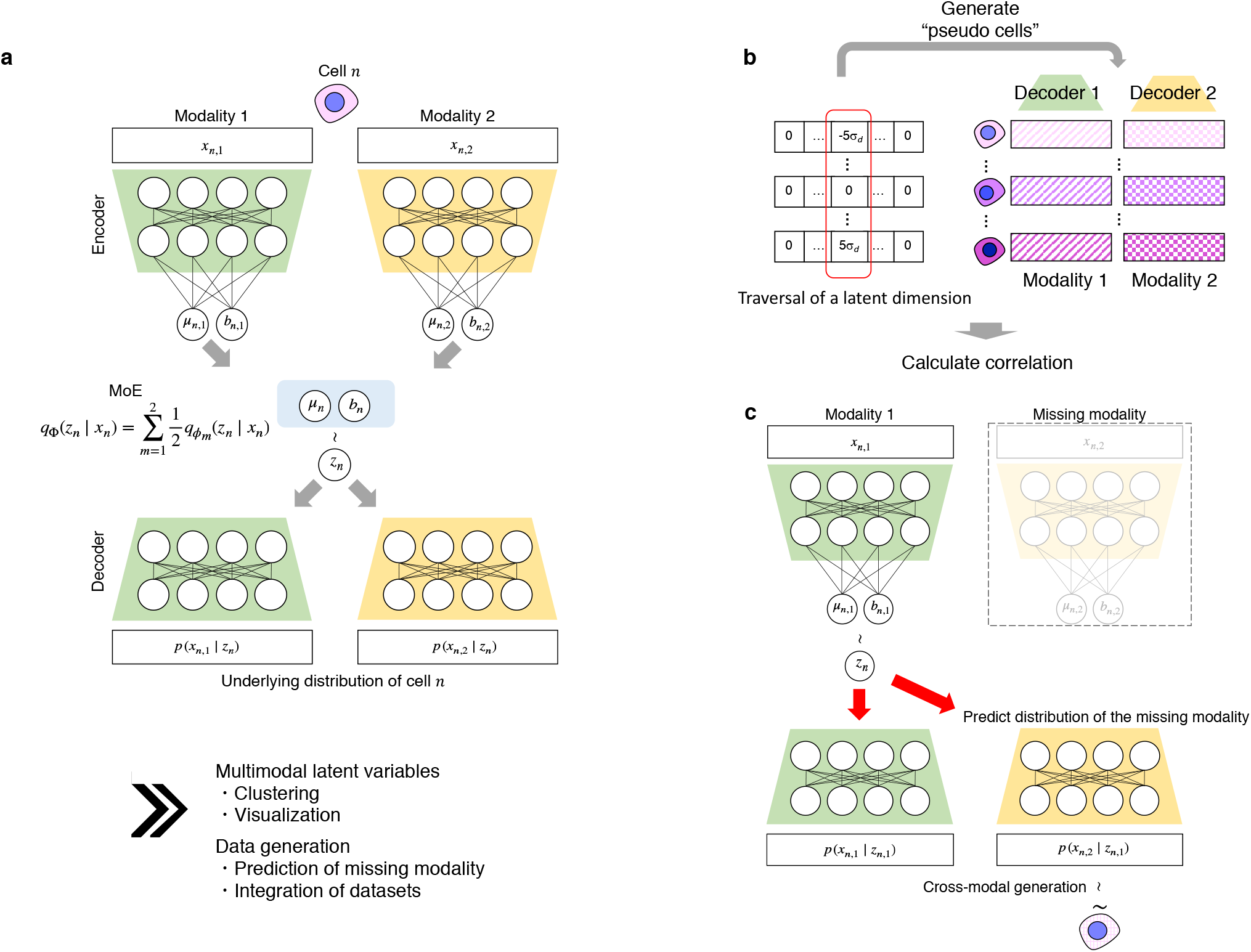
Conceptual view of single-cell multimodal data analysis by scMM. **a**, The underlying model of scMM. scMM takes feature vectors of each modality as input to separate encoders. The model is trained to learn lowdimensional joint variational posterior factorized by a mixture-of-experts (MoE). Decoders reconstruct the underlying probability distributions for data in each modality from latent variables. During the training processes, latent variables from one modality are used to reconstruct data in both modalities. **b**, Schematic view of procedure for finding latent dimension-associated multimodal features by independently traversing each dimension. **c**, Schematic view of cross-modal generation by trained scMM model when one modality is missing. For further details, see the Methods section.

For interpretability, it is helpful to know what features in each modality are associated with each latent dimension. One of the downsides of deep generative models is the difficulty to interpret latent variables compared with linear models, such as the principal component analysis (PCA) [18]. We overcame this limitation by using the generative nature of VAE (Fig. 1b) (see Methods). By sequentially generating pseudocells from different latent values in one dimension with remaining fixed values, we calculated the Spearman correlation for each latent dimension and features in each modality. This enabled the visualization of strongly associated features with each latent dimensions, which can be interpreted as multimodal regulatory programs governing them.

A unique learning procedure of scMM is that encoders are trained to infer latent variables that can reconstruct the probability distributions for not only for its own modality but also for other modalities, as it learns to maximize the expectation 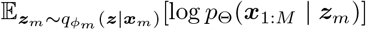 (see Methods). Therefore, the trained scMM model can generate data associated with the missing modality from unimodal single-cell data in both directions, thus achieving cross-modal generation (Fig. 1c). Notably, unlike conventional prediction methods, cross-modal generation by scMM can be performed in both directions across modalities.

### 2.2 scMM extracts biologically meaningful latent variables from single-cell transcriptome and surface protein multimodal data

To validate the performance of scMM in the joint analysis of multimodal single-cell data, we applied our method to a recently published CITE-seq dataset of peripheral blood mononuclear cells (PBMCs) from vaccinated patients, which consisted of the transcriptome and 224 surface protein measurements for over 160,000 cells [5]. In total, 80% of the cells were randomly selected as training data, and the remaining 20% were used as test data. After training the model, all cells were mapped in the latent space and clustering on latent variables was performed with PhenoGraph [19]. Latent variables for each modality and multimodal latent variables were visualized with UMAP (Fig. 2a-c) [20]. To eliminate the possibility of overfitting, we confirmed that the training and test datasets were embedded in shared latent space (Supplementary Fig. 1). Clustering by PhenoGraph discovered 54 cell populations, which can be matched with known cell populations (Fig. 2c). Abbreviations for cell types were as follows: CD4 positive T cell (CD4 T), CD8 positive T cells (CD8 T), gamma-delta T cells (gdT), double-negative T cells (dnT), mucossal associated invariant T cell (MAIT), B cell (B), natural killer cell (NK), CD14 positive monocyte (CD14 Mono), CD16 positive monocyte (CD16 Mono), classical dendritic cell 1 (cDC1), cDC2, plasmacytoid dendritic cell (pDC), hematopoietic stem and progenitor cell (HSPC), erythrocyte (Eryth). Interestingly, compared with the weighted nearest neighbor (WNN) analysis by Seurat, latent variables inferred by scMM separated CD4 and CD8 T into two distinct subgroups (Fig. 1d) [5]. We found that these subgroups have differential expressions of surface proteins which are known to be associated with T cell activation, such as CD30, CD275, and Podoplanin (Fig. 1d, Supplementary Fig. 2a). Furthermore, scMM discovered a clear heterogeneity in CD14 Mono populations that was not revealed by Seurat (Supplementary Fig. 2b). The superior performance of the proposed model may be attributed to the rich expressive power of neural networks used in scMM, whereas the WNN analysis was based on a linear model with limited expressive power. Thus, it was thus unable to capture complex structures in single-cell multimodal data.

**Figure 2:**
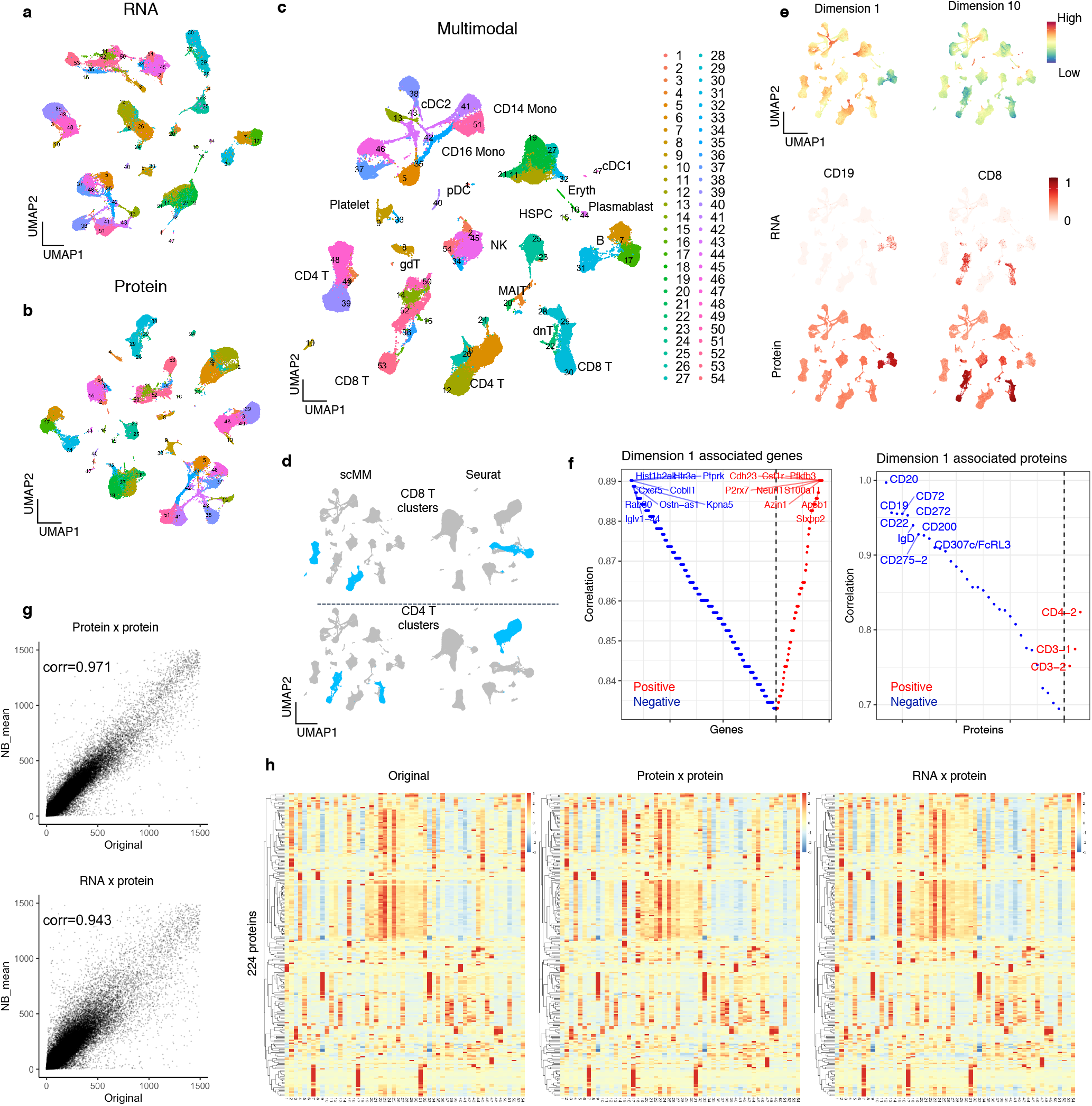
scMM analysis on the PBMC CITE-seq dataset. **a**, **b**, **c**, UMAP visualization of unimodal latent variables for transcriptome, surface protein, and multimodal latentvariables, respectively. Each dot represents a single cell and is color-coded according to the clustering performed on multimodal latent variables. **d**, CD8 and CD4 T cells annotated in scMM and Seurat analysis are color-coded in the UMAP visualization. **e**, Upper panels; Multimodal UMAP visualization colored according to the latent dimension values. Middle and lower panels; UMAP visualization colored according to the transcriptome and protein counts for cell type markers, respectively. **f**, Genes (left panel) and surface proteins (right panel) associated with latent dimension 1. Each feature was aligned based on Spearman correlation coefficient. The y-axis represents the absolute correlation coefficient, and red and blue represent positive and negative correlations, respectively. **g**, NB mean parameters reconstructed from surface protein or transcriptome counts are plotted against original surface protein counts for each cell. Pearson correlation coefficients are shown in the plots. **h**, Heatmap constructed from the original (left panel), unimodal generation (middle panel), and cross-modal generations. Rows and columns represent the measured 224 surface proteins and 54 clusters discovered by PhenoGraph, respectively.

### 2.3 scMM supports result interpretation by providing multimodal features associated with latent dimensions

scMM uses a Laplace prior with different scale values in each dimension, which encourages disentanglement of information by learning axis-aligned representations [11] (see Methods). Visualizing values of latent variables revealed similar patterns with canonical gene and surface protein markers (Fig. 2e). This may indicate axis-aligned encoding of information related to certain cell types. For example, low values in latent dimension 1 were concentrated to B cell and plasmablast clusters (7, 17, 31, and 44) as indicated by the expression of the CD19 gene and protein. This may suggest that latent dimension 1 encodes information on cellular characteristics of B cell and plasmablast. Fig. 2e shows multimodal features strongly associated with latent dimension 1. Genes showing negative correlations include those related to immunoglobulin (Iglv1-44), chemotaxis (Cxcr5), and membrane trafficking (Rab30), and their expressions in B cell and plasmablast clusters were confirmed (Fig. 2e, Supplementary Fig. 3a). Surface proteins are well-known to be associated with the B cell and plasmablast phenotype, such as CD19, CD20, and CD22, which were also detected in negatively correlated features (Fig. 2e, Supplementary Fig. 3b). Collectively, these results validate utilities of interpretable latent representations learned by scMM.

### 2.4 Cross-modal generation by scMM accurately predicts surface protein measurements from transcriptome data

A trained scMM model can generate surface protein measurements conditioned on transcriptome observations (and vice versa, although transcriptome counts are generally acquired simultaneously when measuring surface proteins with sequencing technologies) through cross-modal generation. Using held-out test data, estimates of mean parameters for NB distributions were plotted against original surface protein counts, which exhibited a high correlation not only in transcriptome-to-transcriptome but also in transcriptome-to-protein cross-modal estimation. From estimated NB distributions, surface protein measurements were sampled for each cell, and heatmaps were generated for 54 clusters (Fig. 2h). As a result, a heatmap constructed from cross-modal generation showed a high resemblance to that of the original, confirming cross-modal data generative performance of scMM.

This feature of scMM can be used to predict surface protein measurements from unimodal single-cell datasets, which comprise transcriptome information only. We validated the performance of scMM by predicting surface protein abundance using data from different experimental batches. To compare predicted versus ground-truth, we chose BMNC CITE-seq data, containing approximately 30,000 cells with transcriptome and 25 surface protein information [21]. With scMM trained with the PBMC training data, latent variables were obtained from transcriptome measurements of BMNC data and visualized using UMAP (Fig. 3 a). Hence, BMNC data were successfully embedded in latent space learned from the PBMC training data. Notably, scMM correctly illustrated the enrichment of CD34 positive hematopoietic stem and progenitor cells (HSPC) in the bone marrow, where this population is scarce in peripheral blood (Fig. 3 b, Supplementary Fig. 4 a). In addition, it is noteworthy that scMM embedded CD8 and CD4 T cells in BMNC datasets with activated, CD30 positive T cell subsets found in the PBMC dataset (Supplementary Fig. 4 b-e). This finding is reasonable given that CD30 marks memory T cells, and they reside mainly in the bone marrow [22, 24].

**Figure 3:**
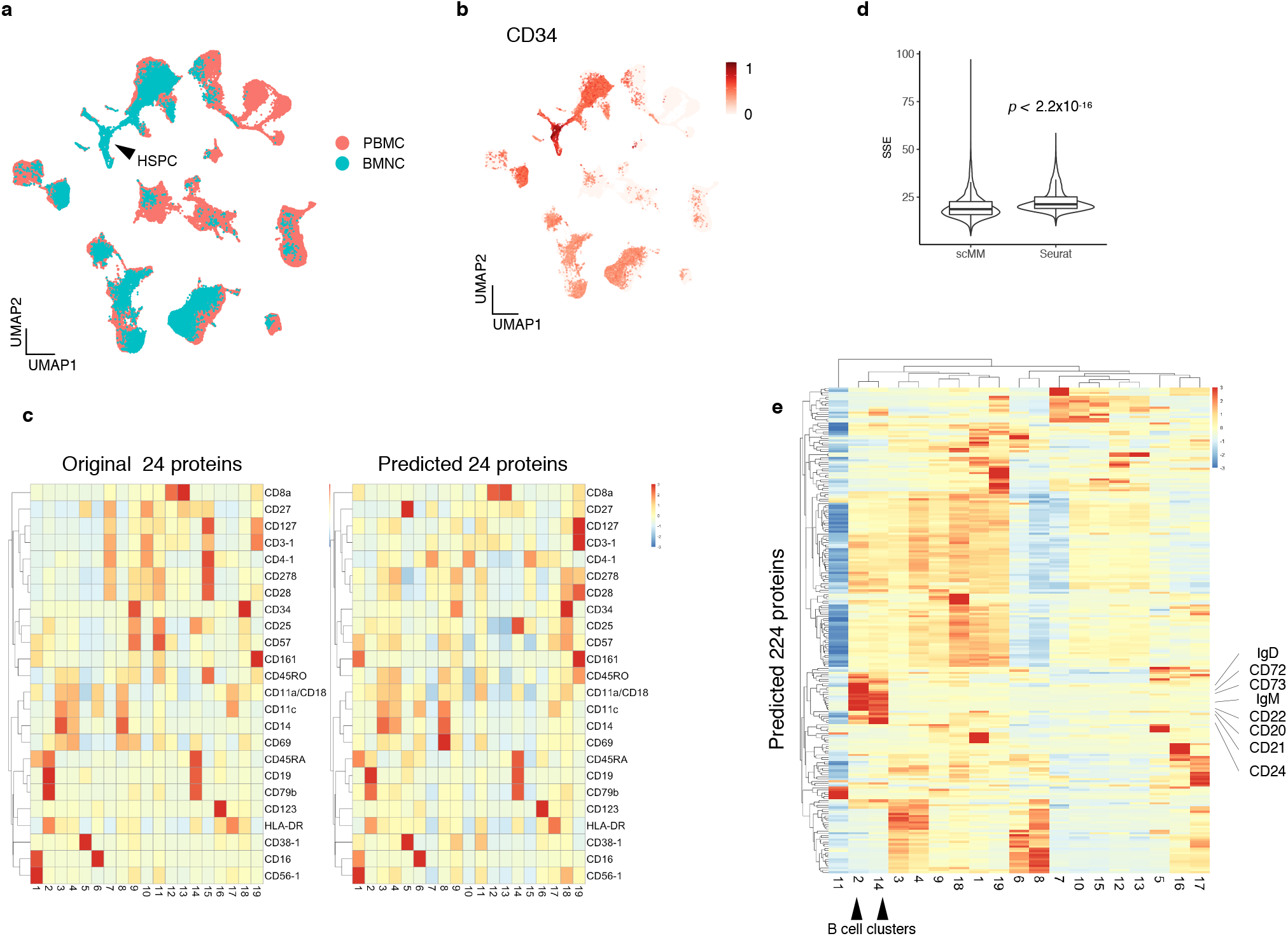
Prediction of surface protein measurements for the BMNC dataset. **a**, Joint UMAP embedding of transcriptome latent variables inferred from PBMC training data and the BMNC dataset. Black arrowhead indicates the HSPC population. **b**, UMAP embedding of the BMNC dataset was colored according to the protein expression levels of CD34. **c**, Heatmaps constructed from original (left panel) and predicted (right panel) surface protein counts. Rows and columns represent 24 shared surface protein markers and clusters discovered by PhenoGraph, respectively. **d**, Benchmarking on surface protein prediction performance of scMM against Seurat. Centered log-transformed data were used to calculate SSEs per cell and plotted for each prediction result. Statistical analysis was performed with the two-sided Wilcoxon signed-rank test. **e**, Heatmap constructed with the predicted 224 surface protein markers. Rows and columns represent surface protein markers and clusters, respectively. Black arrowheads denote B cell clusters, and their markers are indicated.

Subsequently, cross-modal data generation was performed by sampling from NB distributions for surface proteins. Out of 25 surface proteins analyzed in the BMNC dataset, 24 were shared with the PBMC dataset. For 19 clusters discovered by PhenoGraph clustering, expression levels of the shared surface proteins were visualized using a heatmap. The result shows surface protein data generated by scMM well captured the characteristics of the original data (Fig. 3 c). We benchmarked the prediction accuracy of scMM against Seurat, which is currently the state-of-the-art method to predict surface protein from single-cell transcriptome [21]. For comparison, we trained scMM and Seurat using the PBMC training data and used them to predict surface proteins of the BMNC dataset. The sum of squared error (SSE) per cell depicted that the proposed method was more accurate than Seurat in predicting the surface protein (Fig. 3 d). Stochastic sampling processes may be the reason for the higher variation in scMM. Fig. 3 e illustrates all 224 surface proteins predicted by scMM. Notably, prediction by scMM recovered crucial features of cell populations. For instance, B cell clusters (cluster 2 and 14) characterized by CD19 expression in the original data were predicted to have high expression levels of known B cell markers that were missing from the original data (CD72, CD73, CD22, CD20, CD21, CD24, IgD, and IgM) (Fig. 3 e). The trained scMM model can generate surface protein measurements conditioned on transcriptome observations (and vice versa) although transcriptome counts are usually acquired simultaneously when measuring surface proteins with sequencing technologies) through cross-modal generation. Using held-out test data, estimates of mean parameters for NB distributions were plotted against original surface protein counts. The results showed a high correlation not only in transcriptome-to-transcriptome but also in transcriptome-to-protein cross-modal estimations. From these NB distributions, surface protein measurements were sampled for each cell, and heatmaps were generated for 54 clusters (Fig. 2h). Thus, the heatmap for data generated from transcriptome data showed a high resemblance to the heatmap of the original. This confirmed the cross-modal data generative performance of scMM.

This feature of scMM can be used to predict surface protein measurements from unimodal single-cell datasets where only transcriptome information is available. We validated the performance of scMM on the prediction of surface protein abundance using data from different experimental batches. To compare the predicted versus ground-truth values, we chose bone marrow mononuclear (BMNC) CITE-seq data that contained approximately 30,000 cells with transcriptomes and information related to 25 surface proteins [21]. With scMM trained with PBMC training data, latent variables were obtained from transcriptome measurements of BMNC data and visualized with UMAP (Fig. 3a). As a result, BMNC data were successfully embedded in a latent space learned from the PBMC training data. Of note, scMM correctly illustrated the enrichment of CD34 positive HSPC in the bone marrow, wherein this population is very rare in peripheral blood (Fig. 3b, Supplementary Fig. 4a). It is noteworthy that scMM embedded CD8 T and CD4 T in BMNC datasets with activated, CD30 positive T cell subsets found in the PBMC dataset (Supplementary Fig. 4b-e). This is reasonable given that CD30 marks memory T cells, which mainly reside in the bone marrow [22, 24].

Next, cross-modal data generation was performed by sampling NB distributions for surface proteins. Out of 25 surface proteins analyzed in the BMNC dataset, 24 were shared with the PBMC dataset. For 19 clusters discovered by PhenoGraph clustering, expression levels of the shared surface proteins were visualized by heatmaps. The result shows surface protein data generated by scMM captured the characteristics of the original data (Fig. 3c). We benchmarked the prediction accuracy of scMM and compared it with that of Seurat, which is currently the state-of-the-art method for predicting surface proteins from the single-cell transcriptome. For comparison, we trained scMM and Seurat using PBMC training data, and used the trained models to predict surface proteins of BMNC datasets. Sum-of-squared errors (SSEs) per cell showed that our method was more accurate than Seurat’s in predicting surface proteins (Fig. 3d). A higher variation in scMM may be attributed to the stochastic sampling processes. Fig. 3e shows all 224 surface proteins predicted by scMM. Notably, prediction by scMM recovered crucial features of cell populations. For instance, B cell clusters (clusters 2 and 14) characterized by CD19 expressions in the original data were predicted to have high-expression levels of known B cell markers that were missing from the original data (CD72, CD73, CD22, CD20, CD21, CD24, IgD, and IgM) (Fig. 3e).

### 2.5 scMM analysis on single-cell transcriptome and chromatin accessibility multimodal data

Next, we applied scMM on recently reported mouse skin single-cell transcriptome and chromatin accessibility multi-modal data obtained by SHARE-seq [6]. Again, train and test data were obtained by an 80%/20% random split. Latent variables for transcriptome and chromatin accessibility, and multimodal latent variables were visualized by UMAP (Fig. 4a-c). Additionally, embeddings of train and test dataset into shared latent space were confirmed (Supplementary Fig. 5). PhenoGraph clustering on multimodal latent variables showed clusters corresponding to known cell types present in epidermis and hair follicles [24]. Abbreviations for cell types are as follows: cycling interfollicular epidermis (IFE C), basal IFE (IFE B), suprabasal IFE (IFE SB), upper hair follicle (uHF), sebaceous gland (SG), outer bulge (OB), outer root sheath (ORS), companion layer (CP), germinative layer (GL), inner root sheath (IRS), cortex/cuticle (CX), medulla (MED), fibroblast (FIB), dermal sheath (DS), dermal papilla (DP), macrophage (MC), endothelial cell (EC), vascular smooth muscle (vSM), melanocyte (MEL). Visualization of latent variables per dimension discovered similar patterns with certain gene expression levels, thus indicating axis-aligned encoding of information associated with cell types (Fig. 4d). For example, latent dimension 9 seemed to correlate positively with DNA topoisomerase gene Top2a expression levels. Top2a is a marker for cells entering mitosis and is upregulated in proliferative keratinocyte subsets including IFE C, GL, MED, and IRS. By sequentially generating pseudocells while independently traversing latent dimensions, we found genes and peaks strongly associated with the latent dimension 9 (Fig. 4e). Consistent with the cell annotations, genes closely related to the cell cycle, such as Stil, Brca1, and Cdca2, were found in positively associated features. We then sought motif enrichment in detected peaks to reveal latent dimension-associated motifs. Motif enrichment analysis discovered significantly enriched motifs, including FOS::JUNB (*p* = 6.08×10^−21^), TP63 (*p* = 2.86 × 10^−13^), POUF3F3 (*p* = 1.16 × 10^−11^), and MEOX2 (*p* = 1.39 × 10^−4^) (Fig. 4f). By visualizing expression levels and motif scores, we confirmed enrichment of associated genes and motifs in proliferative keratinocyte subsets (Supplementary Fig. 6a, b).

**Figure 4:**
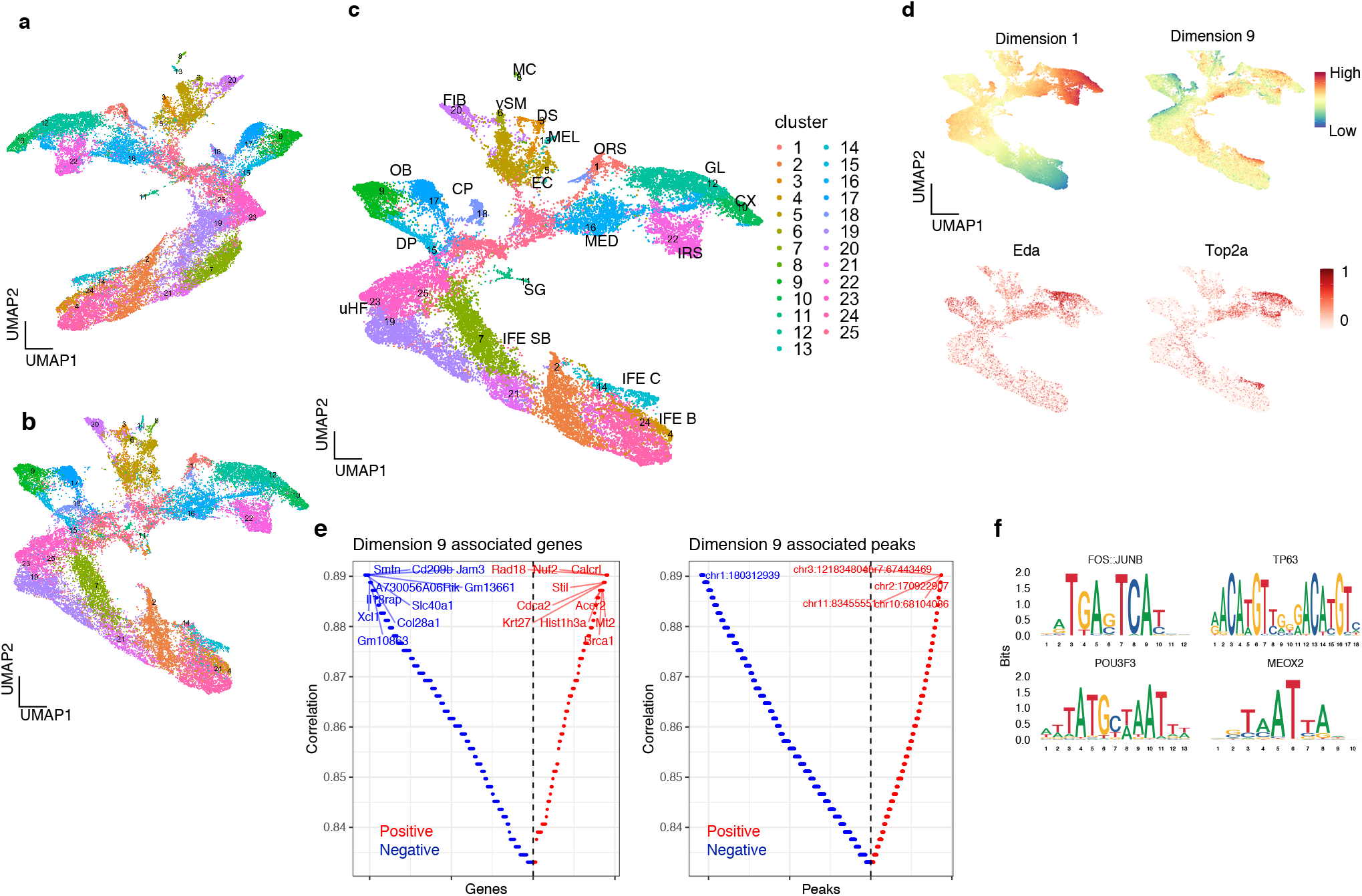
scMM analysis on the mouse skin SHARE-seq dataset. **a**, **b**, **c**, UMAP visualization of unimodal latent variables for transcriptome, chromatin accessibility, and multimodal latent variables, respectively. Each dot represents a single cell and is color-coded according to the clustering performed on multimodal latent variables. **d**, Upper panels: multimodal UMAP visualization color-coded based on latent dimension values. Middle and lower panels: UMAP visualization colored according to transcriptome counts for cell type markers. **e**, Genes (left panel) and peaks (right panel) associated with latent dimension 9. Each feature was aligned based on Spearman correlation coefficient. The y-axis represents the absolute correlation coefficient, where red and blue colors represent positive and negative correlations, respectively. **f**, Motif plot for representative motifs significantly enriched in peaks positively associated with latent dimension 9.

### 2.6 Cross-modal generation of transcriptome measurements contribute to accurate data integration

The predictions of chromatin accessibility from the transcriptome and vice versa are more difficult tasks compared with the prediction of the surface proteins owing to their high dimensionality and sparsity. Specifically, there are only limited methods available for the prediction of chromatin accessibility. A conventional method for predicting the transcriptome from chromatin accessibility is performed by merely summing peak counts within the +2 kb upstream of the gene transcription start site (TSS), which returns the “gene activity matrix (GAM)” [21]. Although GAM corresponds with the transcriptome status of cells to some extent, it accompanies an inevitable loss of information because it ignores the distant interaction of enhancers and TSSs [15].

Regarding the current limitations associated with the prediction of transcriptome and chromatin accessibility from one information set to another, we sought to achieve cross-modal generation by scMM in these modalities. The plot of the estimated means of the NB parameters against the original transcriptome counts showed high correlations in both transcriptome-to-transcriptome and accessibility-to-transcriptome reconstruction (Fig. 5a). Fig. 5b shows heatmaps for 25 clusters on 1126 statistically significant deferentially expressed (DE) genes. The heatmap for cross-modal generation showed similar patterns to those of the original transcriptome data, thus suggesting that generated data captures the characteristics of the original clusters.

**Figure 5:**
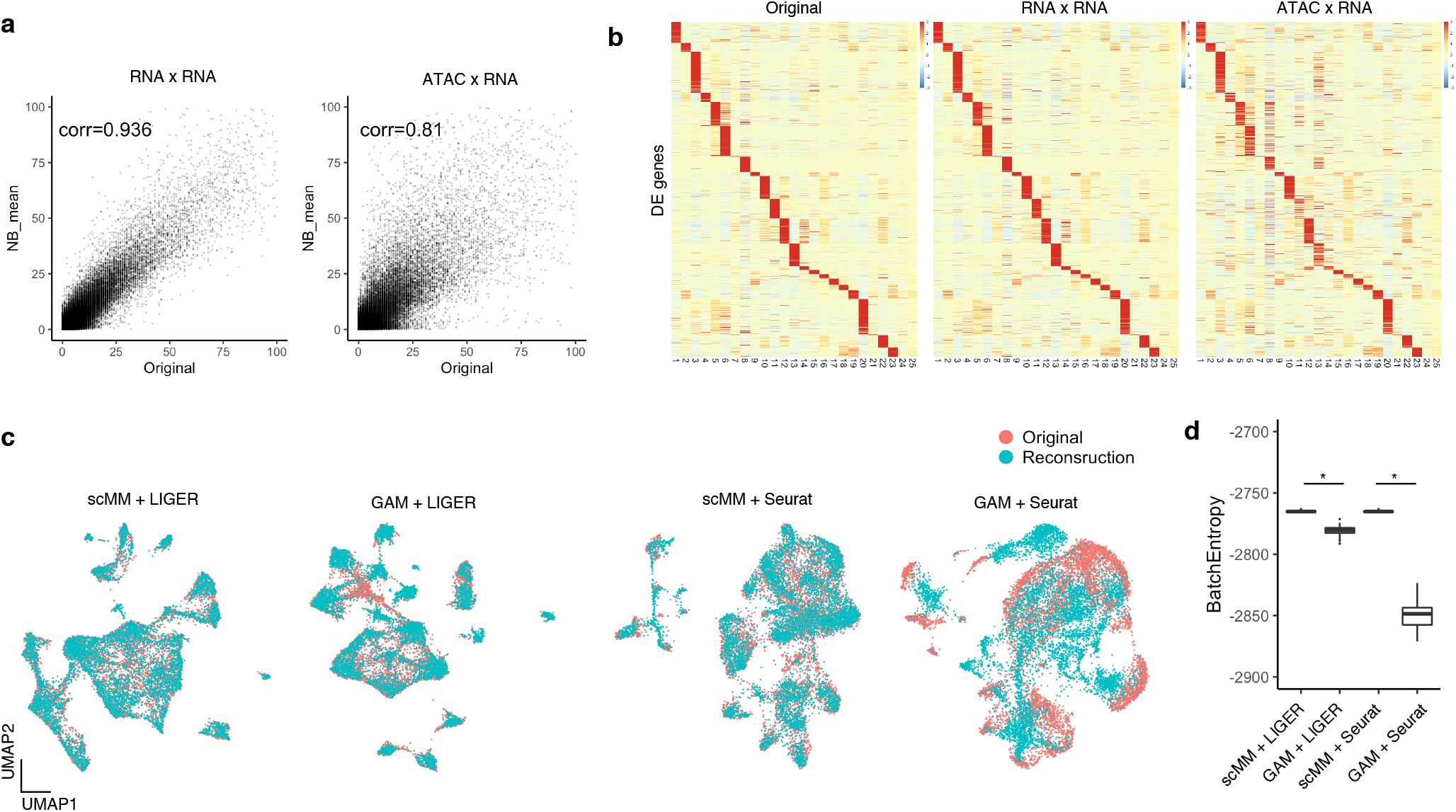
Cross-modal generation from chromatin accessibility to transcriptome leads to better data integration. **a**, NB mean parameters reconstructed from transcriptome or chromatin accessibility counts are plotted against original transcriptome counts for each cell. Pearson correlation coefficients are shown in the plots. **b**, Heatmaps constructed from original, unimodal generation, and cross-modal generation transcriptome data. Rows and columns represent DE genes and clusters discovered by PhenoGraph, respectively. **c**, Joint visualization of original and predicted single-cell transcriptome data integrated by LIGER and Seurat. For the prediction, either scMM or GAM was used. **d**, Boxplot showing entropy of batch mixing for each integration. Statistical test performed with two-sided Wilcoxon rank sum test. * *p* < 2.2 × 10^−22^.

Integration of single-cell data from different modalities is one of the most important goals of modern computational biology. Recently developed single-cell integration tools, including LIGER and Seurat, require the conversion of chromatin accessibility to transcriptome by creating GAM to perform integration [21, 25]. Recent research using single-cell multimodal data as ground-truth has reported that this approach often fails to identify corresponding cells correctly [15]. We reasoned that the use of cross-modal generated transcriptome data by scMM may lead to more accurate integration as it considers all chromatin sites upon prediction. First, we obtained predicted transcriptome measurements for each cell in test data by either cross-modal generation of scMM or by the construction of GAM. Then, we integrated predicted and original single-cell transcriptome data into shared space by LIGER and Seurat. Quantitative evaluation of the integration was performed by calculating the entropy of batch mixing, which is a measurement of how well samples from two batches are integrated, where a higher entropy indicates better integration [26]. With both LIGER and Seurat, integration of cells generated by scMM resulted in better embedding with original cells compared with those performed with GAM (Fig. 5c, d). Collectively, these results suggest that scMM has a promising potential in generating transcriptome data that precisely reflect chromatin accessibility, and significantly contributes to single-cell integration analysis when used in combination with existing methods.

### 2.7 scMM achieves chromatin accessibility prediction

Next, we investigated the prediction of chromatin accessibility from the transcriptome. In contrast to the transcriptome measurements where counts continue to increase with the abundance of mRNA in each cell, there are theoretically only two states in the chromatin accessibility: open or closed. Therefore, larger peak counts only reflect sequences with favorable Tn5 binding, or they are just random events. Thus, the prediction model is required to discriminate zero and nonzero values, rather than predict the absolute counts. Fig. 6a shows the estimated mean parameters of the ZINB distribution for held-out test datasets against the original peak counts. Generally, estimated mean parameters were lower than the original peak counts, which may reflect the low detection rate of open chromatin. Of note, for the peaks with zeros of the original count, the mean ZINB parameter concentrated at zero, thus indicating that scMM accurately captured the closed chromatin (Fig. 6a). Unimodal and cross-modal generation of chromatin accessibility measurements for test datasets was performed by sampling the estimated ZINB distributions. Interestingly, 4018 statistically significant deferentially accessible (DA) heatmap peaks showed high similarities for cross-modal generation and original data (Fig. 6b). In addition, Motif scores in original clusters were accurately recovered by cross-modal generation (Fig. 6c). To investigate the cross-modal generated chromatin accessibility data, we analyzed coverage peaks in the Lef1 and Krt1 gene regions, which are essential markers for anagen hair follicle keratinocytes and permanent epidermis keratinocytes, respectively. Coverage plots showed chromatin accessibility data generated by scMM reconstructed peaks specifically detected in the keratinocyte subsets, further confirming the cross-modal data generation performance of scMM (Fig. 6d). Notably, cross-modal generation by sampling from predicted ZINB distributions allows the formulation of sparse representations of high-dimensional chromatin accessibility data that is memory-efficient compared with dense representations. For 6955 cells in the test dataset, memory sizes were 504 MB versus 10.6 GB for sparse (sampled peak counts) and dense (ZINB mean parameters) representations, respectively.

**Figure 6:**
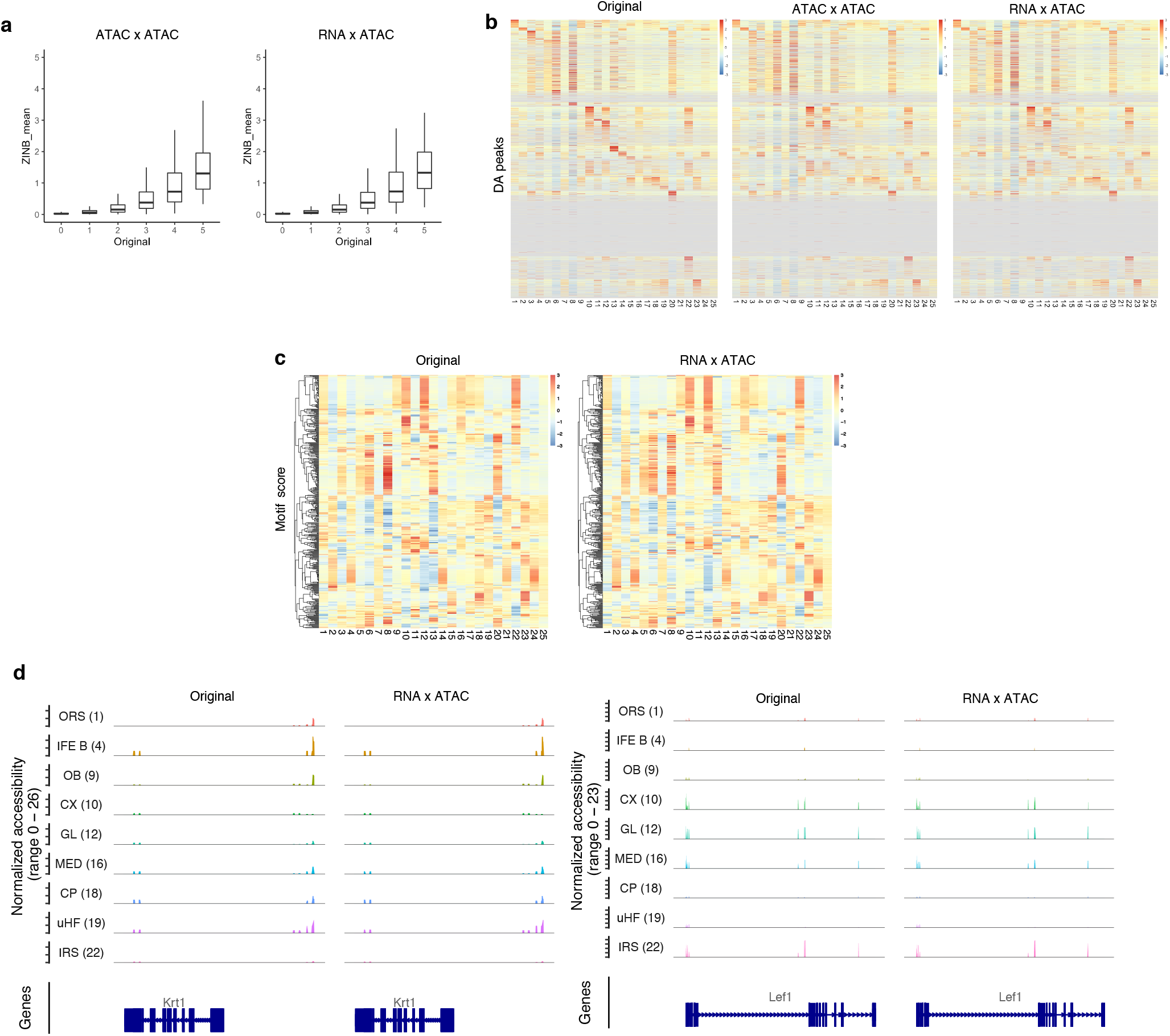
scMM accurately predicts chromatin accessibility from transcriptome data. **a**, ZINB mean parameters reconstructed from transcriptome or chromatin accessibility counts are plotted against original chromatin accessibility counts for each cell. 25 million data points were randomly selected for plotting. **b**, Heatmaps constructed from original, unimodal generation, and cross-modal generation chromatin accessibility data. Rows and columns represent DA peaks and clusters discovered by PhenoGraph, respectively. **c**, Heatmaps of motif scores for original and cross-modal generation data. Rows and columns represent motif scores and clusters, respectively. **d**, Coverage plots for representative clusters within gene regions of Lef1 and Krt1.

## 3 Discussion

The rapidly evolving field of single-cell multimodal analysis requires a method for joint analysis of the obtained data. scMM was designed to meet this demand. In this report, we showed that scMM extracts low-dimensional latent variables from multimodal single-cell data that are useful for downstream analysis, such as clustering. In addition, we leveraged the data generative nature of scMM to compensate for the difficulties in interpreting deep generative models. Exploring the multimodal regulatory programs associated with each latent dimension is expected to provide deeper insights into the clusters discovered in scMM analyses.

Furthermore, we demonstrated that cross-modal generation with scMM achieved accurate prediction of measurements in different modalities. Also, these predictions can be used for integrating multiple unimodal datasets. Benchmarking of scMM against the state-of-the-art prediction tool and conventional integration approaches proved the superiority of scMM in these tasks. These features of scMM will lead to the effective utilization of the accumulating unimodal single-cell databases that are annotated and well-characterized. As a limitation of scMM, however, it may be challenging to generate cell populations that are significantly different from those used to train the model. It is expected that as the construction of large-scale multimodal single-cell atlas progresses, more training data will become available, which would mitigate the problem [27].

One of the strengths of scMM is its extensibility, as it can be applied to any modality by constructing the model with different distributions. In addition to the modalities dealt in this study, applications to other multimodal data, such as the single-cell transcriptome and DNA methylome, constitute attractive future directions [28]. Extending scMM to several multimodal single-cell data may decipher novel cellular states or functions regarding transcriptome, epigenome, and proteome. Application to single-cell data with spatial information would be an exciting research question because, unlike other modalities, coordinates in spatial data is meaningful only when the relative positional relationships with other cells are considered [29]. In essence, we expect that the proposed model will be the foundation for deep generative models for multimodal single-cell data from the scope of interpretable latent feature extraction and cross-modal generations.

## 4 Methods

### 4.1 Mixture-of-experts multimodal VAE model for single-cell multiomics data

This section briefly describes MoE multimodal VAE (MMVAE), which scMM is based on. Further mathematical details on MMVAE are described in the original publication [11]. Suppose there is single-cell multimodal dataset ***x***_1:*M*_. For current multimodal single-cell data, *M* = 2 in general. MMVAE aims to learn multimodal generative model 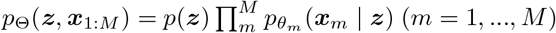, where 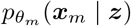 represents the likelihood for each of *m*th modality parameterized by the decoder’s deep neural network. The training objective is to maximize the marginal likelihood *p*_Θ_(***x***_1:*M*_), which is approximated by optimizing the evidence lower bound (ELBO) by stochastic gradient descent (SGD). As shown in Eq. 1, formulation of the ELBO requires approximation of the true posterior by joint variational posterior *q*_Φ_(***z*** | ***x***_1:*M*_).

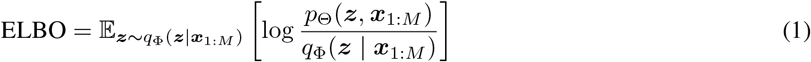

The key idea of MMVAE is to factorize the joint variational posterior with a MoE; 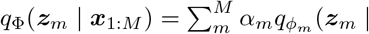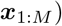, *α*_*m*_ = 1/*M*, where 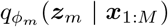 is the variational posterior for the *m*-th modality parameterized by the encoder deep neural network. Using stratified sampling [32], the ELBO can be formulated as:

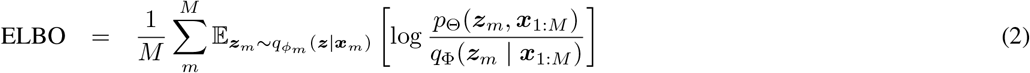

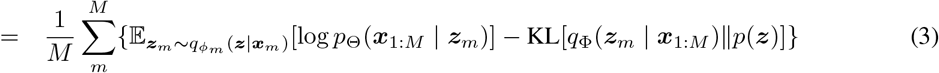

Intuitively, the first term could be interpreted as the goodness of reconstruction for all *M* modalities from latent variables for the *m*th modality. The second term regularizes the model so that variational posterior follows the prior distribution.

For the prior *p*(***z***), we used the Laplacian distribution with a zero mean and a scaling constraint 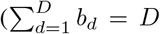, wherein *b*_*d*_ is a scale parameter for the *d*th dimension, and *D* is the number of latent dimensions) [11]. Scale parameters for prior were learned from data through SGD.

Likelihoods 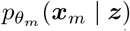 were selected according to the data distribution characteristics of each single-cell modality. NB distribution was used for transcriptome and surface protein counts, and ZINB distribution was used for chromatin accessibility peak counts.

For all modalities, row counts were normalized by dividing with the sequencing depth of each cell, multiplying with the scale factor (10,000) and used as input to encoders. Mean parameters estimated by decoders were processed with reverse processes, and log likelihoods were calculated with raw count data.

### 4.2 Model architecture and optimization

Optimization was performed with an Adam optimizer with AMSGrad [30]. Hyperparameter optimization was performed by Optuna [31]. For CITE-seq data, three hidden layers with 200 hidden units were used for both modalities. For SHARE-seq data, two hidden layers with 500 units for transcriptome and 100 hidden units for chromatin accessibility were used. Learning rates were set to 2e-3 and 1e-4 for CITE-seq and SHARE-seq data, respectively. We used deterministic warm-up learning scheme for 10 and 50 epochs, with maximum of 20 and 100 epochs for CITE-seq and SHARE-seq data, respectively [33]. After deterministic warm-up, early stopping with a tolerance of 10 epochs was applied. We observed that minor changes in hyperparameters did not significantly affect the analyzed results.

### 4.3 Data preproccessing

For transcriptome count data, 5000 most variable genes were first selected by applying the Seurat *FindMostVariable* function to log-normalized counts. Raw counts were used for model input. For chromatin accessibility data, the top 25% peaks were selected for input using Seurat’s *FindTopFeatures* function. No preprocessing and feature selection were performed on surface protein count data.

### 4.4 Cluster analysis

Clustering was performed with the R package Rphenograph with nearest neighbor numbers *k* = 20 and *k* = 15 for human PBMC/BMNC CITE-seq data and mouse skin SHARE-seq data, respectively.

For CITE-seq data, cell types were manually annotated with known surface protein markers and by referring to the original report [5]. For SHARE-seq data, manual annotations were performed with the mouse skin single-cell data portal http://kasperlab.org/mouseskin.

Heatmaps were generated with the R package pheatmap. For gene, surface protein, and chromatin accessibility, z-scores for total feature counts normalized by the total sequencing depth per cluster were used to generate heatmaps. For motif scores, z-scores for median values were used.

### 4.5 Visualization of latent representations

Mean parameters for variational posteriors in each modality and MoE were used as latent variables. Latent variables obtained from trained models were visualized on the two-dimensional space using the “umap” package in R.

### 4.6 Detection of latent dimension-associated features

We generated series of pseudocells by independently traversing latent dimensions. This approach was inspired by the study on the original MMVAE paper [11]. Using the learned standard deviation for the *d*th dimension *σ*_*d*_, with other dimensions fixed to zero, we linearly changed the *d*th dimension from −5*σ*_*d*_ to 5*σ*_*d*_ at a rate of 0.5*σ*_*d*_. The obtained latent vectors were then decoded for *M* modalities and resulted in 20 pseudocells. Spearman’s correlation coefficients were calculated for traversed latent dimensions and features in each modality. Latent dimension-associated features were selected using p value thresholds that produced a reasonable number of associated features, namely, *p* < 1 × 10^−12^, *p* < 1 × 10^−3^, and *p* < 1 × 10^−21^ for genes, proteins, and peaks, respectively.

### 4.7 Motif analysis

Motifs enriched in latent dimension-associated peaks were obtained by the *FindMotifs* function in Signac [34]. Motif scores were calculated using the chromVAR wrapper function *RunChromVAR* in Signac [35].

### 4.8 Benchmarking of scMM

Surface protein prediction by Seurat was performed following the official tutorial. Specifically, anchors between PBMC training data and BMNC data were calculated, and prediction was performed with the *MapQuery* function. Given that Seurat returns the centered log-transformed measurements, the predicted results of scMM were also transformed to compare SSEs per cell. GAM was generated by the *GeneActivity* function of Signac. Cross-modal transcriptome reconstruction by scMM and GAM were integrated with original data by LIGER and Seurat following official tutorials with default parameters. The entropy of batch mixing was calculated as described in a previous study [26]. Briefly, for 100 randomly chosen cells, their 100 nearest neighbors were used to calculate the batch proportion *x*_*i*_, where *x*_1_ + *x*_2_ = 1. Regional entropy was estimated according to *E* = *x*_1_ log *x*_1_ + *x*_2_ log *x*_2_, and entropy of batch mixing was calculated as their sum. For boxplot, this process was repeated 100 times with different randomly chosen cells.

### 4.9 Statistical analysis

The function *findMarkers* of the R package scran was applied on log-normalized gene counts to detect DE genes (p value < 0.05, FDR < 0.1). For detection of DA peaks, the function *FindMarkers* in the R package Signac was used with a logistic regression mode (p value < 0.05, log2-fold change > 0.5). Wilcoxon signed-rank test and sum rank test were performed with the function *wilcox.exact* in the R package exactRankTests.

### 4.10 Data availability

PBMC and BMNC CITE-seq data are available at the official website of Seurat https://satijalab.org/seurat/. Mouse skin SHARE-seq data is available under the accession number GSE140203. The scMM model was implemented with Python using PyTorch deep learning library, and code is available at https://github.com/kodaim1115/scMM.

## 5 Acknowledgements

Not applicable.

## 6 Author contributions

K.M. designed and performed experiments under supervision of H.N. and T.S. All authors read and approved the final manuscript.

