## Supplementary Fig. for "scMM: Mixture-of-experts multimodal deep generative model for single-cell multiomics data analysis"

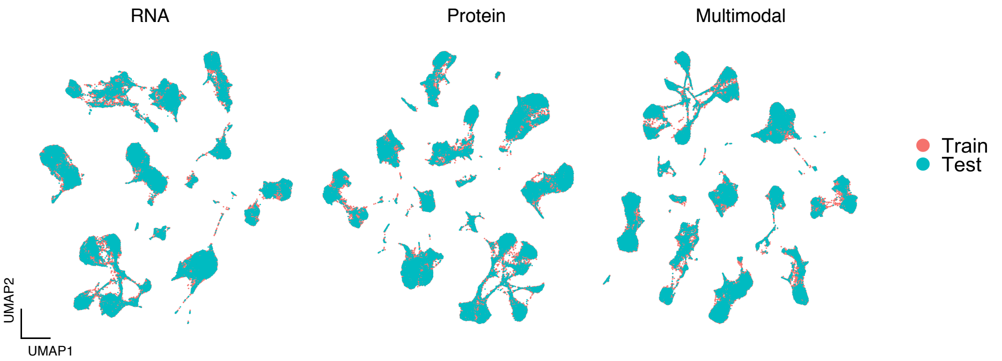


**Supplementary figure 1. scMM embeds the train and test PBMC CITE-seq datasets into shared latent space.** UMAP visualization of unimodal and multimodal latent variables color-coded by assignments to train or test dataset.

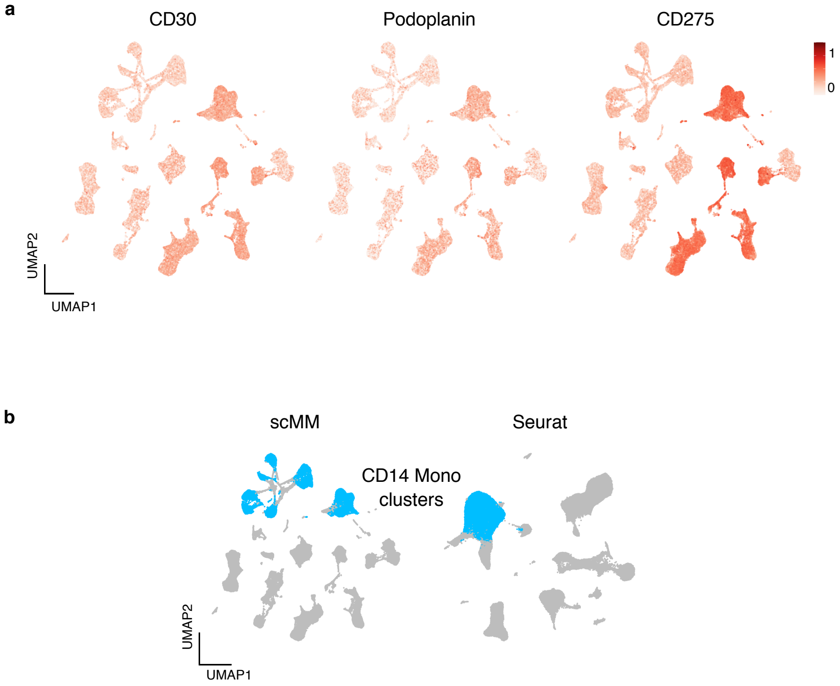


**Supplementary figure 2. scMM revealed previously undiscovered heterogeneity in the PBMC dataset. a**, UMAP visualization of multimodal latent variables colored according to the expression levels of surface proteins differentially expressed in subsets of CD8 and CD4 T populations. **b**, CD14 Mono populations annotated in scMM and Seurat clustering analysis are indicated.

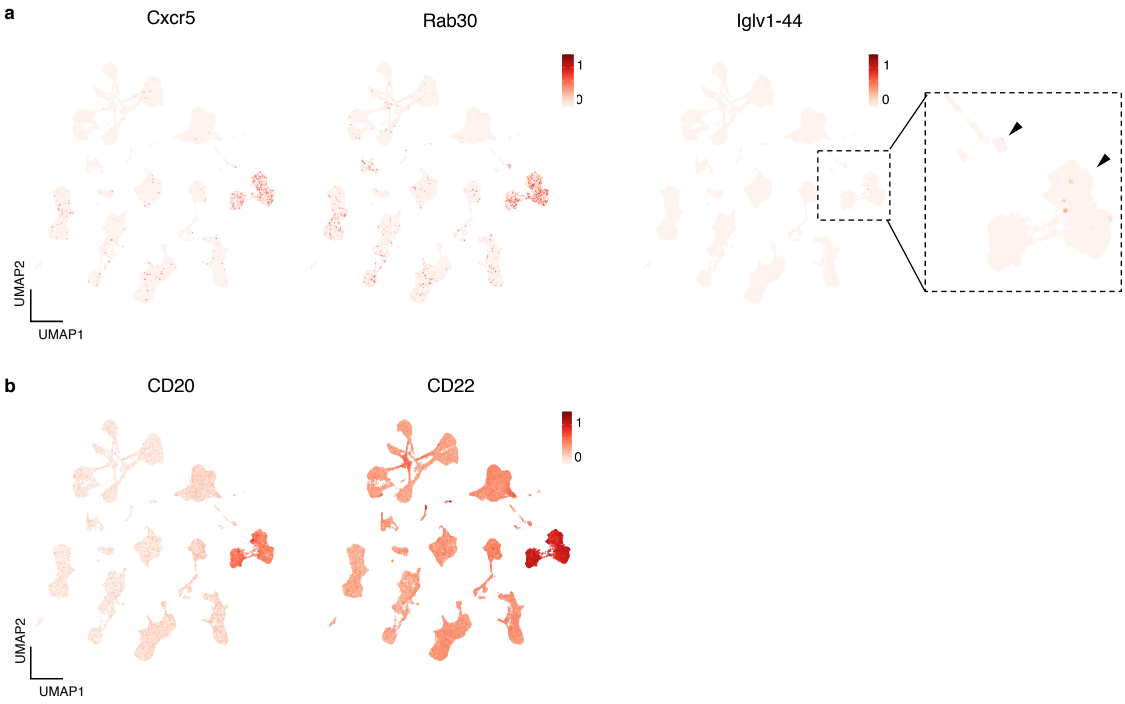


**Supplementary figure 3. Genes and surface proteins detected to be associated with the latent dimension 1 enriches in B cell and plasmablast populations. a**, UMAP visualization of multimodal latent variables colored by expression levels of Cxcr5, Rab30, and Iglv1-44. Black arrowheads in enlarged panel indicate expression of Iglv-44. **b**, UMAP visualization of multimodal latent variables colored according to the expression levels of CD20 and CD22.

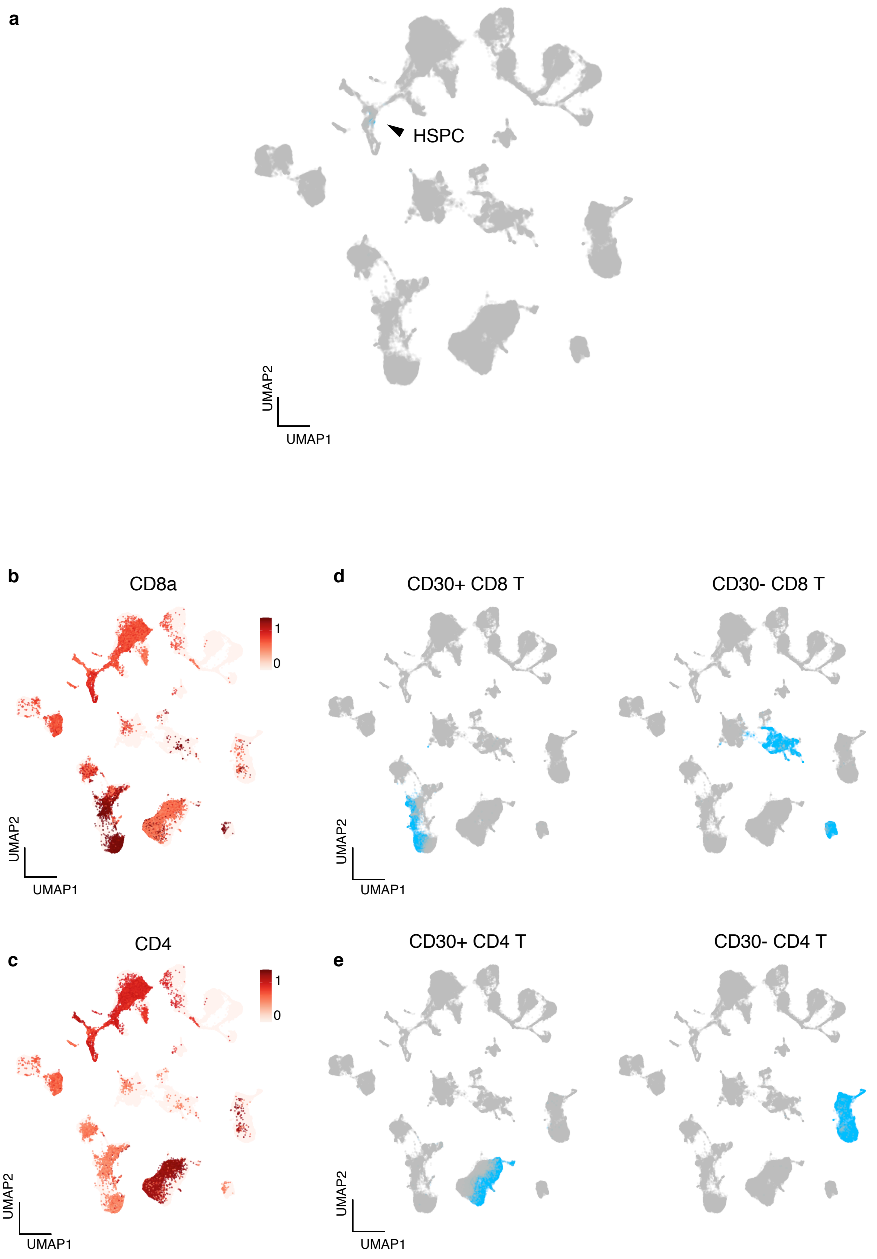


**Supplementary figure 4. Embeddings of BMNC cells into the latent space by scMM captures the biological properties of bone marrow populations. a**, Joint UMAP visualization of BMNC and PBMC transcriptome unimodal latent variables. HSPC population annotated in PBMC dataset is color-coded and indicated by black arrowhead. **b**, **c**, UMAP embeddings for BMNC dataset is colored according to the expression levels of CD8 and CD4, respectively. **d**, **e**, CD8 and CD4 subsets in the PBMC dataset with and without CD30 expression are color-coded.

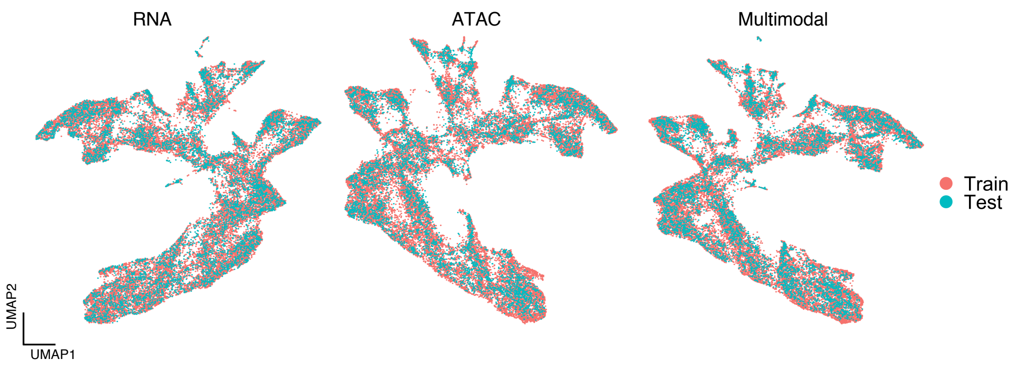


**Supplementary figure 5. scMM embeds train and test mouse skin SHARE-seq datasets into the shared latent space.** UMAP visualization of unimodal and multimodal latent variables color-coded by assignments to either train or test dataset.

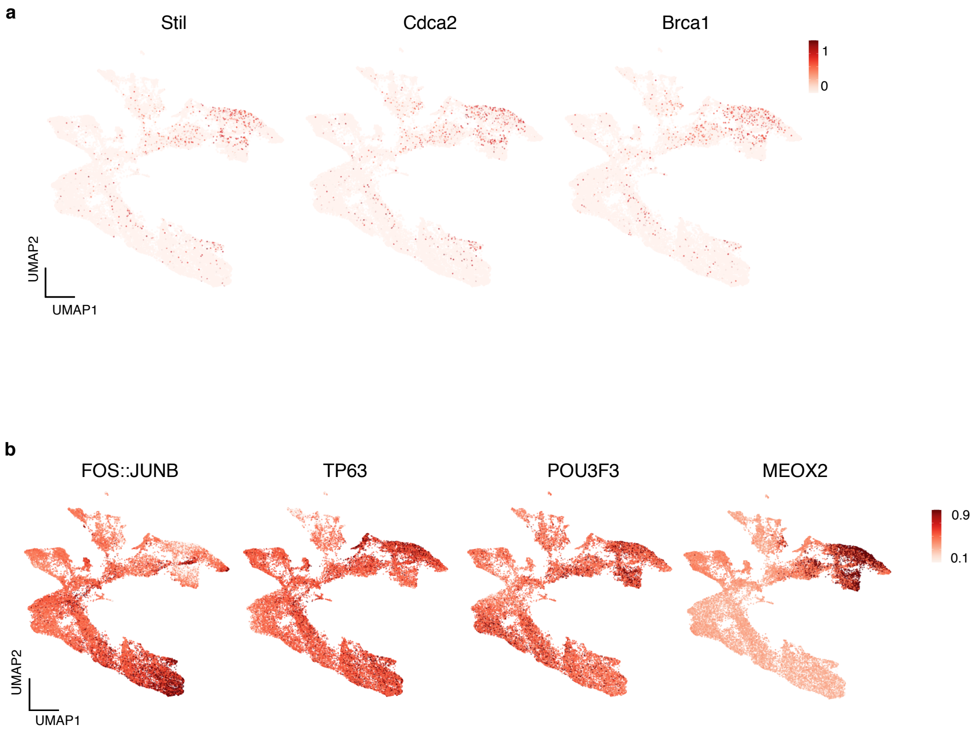


**Supplementary figure 6. Gene and motifs detected to be associated with latent dimension 9 enriches in proliferative keratinocyte subsets. a**, UMAP visualization of multimodal latent variables colored according to the expression levels of Stil, Cdca2, Brca1. **b**, UMAP visualization of multimodal latent variables colored according to the motif scores for FOS::JUNB, TP63, POU3F3, and MEOX2.
